# When tension is exciting: an EEG exploration of excitement in music

**DOI:** 10.1101/637983

**Authors:** Amelia Turrell, Andrea R Halpern, Amir-Homayoun Javadi

**Affiliations:** School of Psychology, University of Kent, Canterbury, UK; Department of Psychology, Bucknell University, Lewisburg, Pennsylvania, USA; Institute of Behavioural Neuroscience, Department of Experimental Psychology, University College London, London, United Kingdom; School of Rehabilitation, Tehran University of Medical Sciences, Tehran, Iran

**Keywords:** Music, Drop, EEG, Emotion, Excitement, Expectancy, Anticipation, Tension

## Abstract

Music powerfully affects people’s emotions. In particular, moments of tension and deviation in musical features, including frequency, pitch, and rhythm (known as a Drop), are associated with positive emotions. However, the neuro-correlates of Drops emotive effects have never been explored. Thirty-six participants listened to music pieces containing a Drop, while undergoing continuous EEG, and rated felt excitement. Source reconstruction of EEG data showed significantly different activity in five brain regions before and after Drops: pre- and post-central gyri (PreCG and PostCG), and precuneus (PCUN) were more active before Drops and the inferior frontal gyrus (IFG), and middle frontal gyrus (MFG) were more active after Drops. Importantly, activity in the IFG and MFG showed a strong correlation with subjective excitement ratings during Drop apprehension. These results suggest expectancy is important to the induction of musical emotions, in agreement with the ITPRA theory. Specifically, when Drops are expected but do not occur immediately, moderate tension is induced. Strong positive emotions then ensue when expected deviations finally occur, due to contrastive valence. This is reflected in significant brain activity for regions associated with high arousing, pleasurable emotions, such as excitement.

## 1 Introduction

Music can produce reliable and powerful emotional responses in listeners, which can increase music engagement (Habibi & Damasio, 2014; Menon & Levitin, 2005; Särkämö & Soto, 2012). While there are numerous studies showing different brain areas associated with passive music apprehension and processing musical features, such as rhythm, pitch, and melodies (Habibi & Damasio, 2014; Särkämö & Soto, 2012; Meltzer et al., 2015; Rolison & Edworthy, 2013) little research has explored the specific effects of highly anticipated but deviant music moments on brain activity and emotions. Different music genres create these deviants in different ways, inducing contrasting levels of tension resulting in different levels of anticipation and associated emotions (Koelsch, 2014). In this study, we investigated effects of these sudden changes on prefrontal cortex (PFC) brain activity and positive emotions, including excitement, taking advantage of a dance music technique in which tension and deviants are created; referred to as Drops. These are points of dramatic variation in the pace, tempo, beat, bass, volume, tone, frequency and/or rhythm of a song. The music builds in these features and abruptly stops or slows to then change.

Three brain areas are consistently shown to be involved in processing musical features during passive music listening, such as structure, rhythm, pitch, and melodies: the inferior frontal gyrus (IFG), dorsolateral prefrontal cortex (DLPFC), and the middle frontal gyrus (MFG) (Kunert, Willems, Casasanto, Patel, & Hagoort, 2015; Lappe, Steinsträter, & Pantev, 2013; Nan & Friederici, 2013; Royal, Zendel, Desjardins, Robitaille, & Peretz, 2018; Thaut, Trimarchi, & Davide, 2014). These areas are also associated with experiencing positive music emotions including; pleasantness, anticipation, empathy, and excitement, as well as physiological responses indicative of positive feelings, such as ‘chills’ (Greene, Flannery, & Soto, 2014; Koelsch, Fritz, Cramon, Müller, & Friederici, 2006; Lehne, Rohrmeier, & Koelsch, 2014; Tabatabaie et al., 2014; Wallmark, Deblieck, & Iacoboni, 2018).

Because music unfolds over time, two musical mechanisms of tension and deviation have previously been associated with positive musical emotions and the previously mentioned brain activity. Perceived music emotions relate to the syntactic structure and experience of musical features, as well as to their predicted and expected future structure; both in terms of anticipation or tension of what is going to happen and of novel, unexpected deviations (Jones, 1982; Huron, 2006; Koelsch, 2014; Kunert et al., 2015; Vuust & Witek, 2014). According to Huron’s (2006) ITPRA theory (imagination, tension, prediction, reaction and appraisal), suggesting highly anticipated music moments create tension enabling greater positive emotions when predicted deviations occur due to contrastive valence (Huron, 2006). Contrastive valence refers to the phenomenon where pleasant emotions are increased when a positive stimuli response immediately follows a negative response (Huron, 2006).

Research has shown music containing high anticipation and tension for predicted or unexpected alterations in its continuous framework evoke pleasant emotions, such as excitement, satisfaction, and surprise despite prior knowledge of the musical piece (Arjmand, Hohagen, Paton, & Rickard, 2017; Koelsch, 2014; Koelsch, Fritz, & Schlaug, 2008; Omigie, 2016; Zentner, Grandjean, & Scherer, 2008). Musical emotion associated with anticipation and deviation has been linked to increased activity in the same three brain areas mentioned above (Bianco et al., 2016; Lehne et al., 2014; Seger et al., 2013), as well as to experiencing pleasant musical emotions (Daly et al., 2014; Lehne et al., 2014; Tabei, 2015; Wallmark et al., 2018).

Here, we explored the relationship between positive emotions induced by Drops and brain activity (EEG and source reconstruction). Specifically, the present research focused on brain activity related to subjective excitement ratings to a 20 second section of a music including Drops varying in strength. We recorded EEG during the presentation of song clips which were used to develop 3D source reconstructions of brain activity across Drop apprehension. We also recorded excitement ratings after each clips. These ratings were used to look at parametric modulation of the brain activity. We hypothesised that brain regions including the IFG, DLPFC, and MFG would increase in activity in response to Drop processing, and that this activity would correlate with subjective ratings of excitement.

## 2 Method

### 2.1 Participants

A total of 36 participants took part in the study (24 females, age mean [SD] = 20.360 [4.372]). Participants were required to have normal or corrected-to-normal vision and hearing. Participants gave written informed consent. The protocol of the study was approved by the local ethics committee at the University of Kent.

### 2.2 Stimuli

Originally, 180 song clips that contained a Drop were extracted from a range of electronic dance music (EDM) genres, including: Dance, Drum & Bass, Dub-Step, House, and Trance. Such music is created electronically with devices such as synthesisers and computers, for the purpose of dancing and is conventionally played in clubs, parties, and raves (Solberg & Jensenius, 2017; Thompson & Stevenson, 2015).The sub genres follow a similar overall structure including structured versus and chorus’ containing Drops, yet differ generally in tempo, frequencies and Drop strengths. Each song clip was 18-22s containing a Drop with 1s fade-in and -out. Pre-Drop consisted of 14s with a jitter of ±2s uniform distribution. Post-Drop was 4s. Undergraduate students (N = 156 in three groups) rated the familiarity, strength, and experienced excitement for each Drop using a 5-point sale (1 = ‘Not familiar at all’ and 5 = ‘Extremely familiar’) and 10-point scale with 10 being ‘Extremely Strong/Exciting’ and 1 being ‘Not strong/exciting at all’, respectively. Highly familiar songs were then excluded and 90 song clips varying in Drop strength from remaining songs were randomly selected to create the stimulus set for the main study.

### 2.3 Procedure

Each trial began with a fixation cross of 3s jittered for ±1s with uniform distribution. Subsequently, a song clip was presented alongside a fixation point on which participants were asked to keep their gaze, minimizing eye movement. Following the song clip, a question mark was displayed. The participants’ task was to indicate how they felt when listening to the song on a 9-point scale, with 9 being ‘Very Exciting’ and 1 being ‘Not Exciting’ via the computer keyboard. Participants were asked to respond as quickly as possible within a 2 second period.

### 2.4 EEG Recording

EEG was recorded continuously from 32 Ag/AgCl electrodes with a BrainVision QuickAmp-72 amplifier system (Brain Products, Germany) placed according to the 10 – 20 electrode placement system. Raw EEG was sampled at 512 Hz with 12-bit resolution. One electrode was used as Ground and a further two around the right eye to record vertical and horizontal eye movements.

### 2.5 EEG Analysis

Our interest was to infer neural substrate of excitement in response to Drops. Therefore, the analysis was done at the source level. We used 3D source reconstruction to have an indication of the location of the activity, and to look at average activity over a period before and after the Drops.

EEG data were analysed using SPM v12 (statistical parametric mapping, Wellcome Trust, London, UK). Data was filtered for 0.5-48Hz using 7th order Butterworth filter, montaged based on average electrode activity, and downsampled to 128Hz data. Then, eye-blinks were removed using activity of the FP2 electrode. Spatial confounds were indicated based on Singular Value Decomposition (SVD) mode and sensor data was corrected using Signal-Space Projection (SSP) correction mode. A maximum of two components of spatial confounds was removed from the EEG data. At the next stage, 1 second pre-Drop and 1 second post-Drop epochs were extracted. Finally, an automatic artefact detection algorithm was applied to the epochs, rejecting those with more than 100mv peak to peak amplitude. A 3D source reconstruction algorithm was then used to extract brain maps of activity sources. EEG Boundary Element Methods (BEM) with normal mesh resolution was used on the SPM template and cortical smoothing for eight iterations. Therefore, 2×90 brain volumes were created for the pre- and post-Drop events. Two analyses were conducted:

Absolute activity – in the first level analysis, contrast images were created using paired-sample t-tests on each pair of pre- and post-Drop. In the second level analysis, these contrast images were subjected to a one-sample t-test.

Parametric modulation – subjective ratings were used to examine the modulation of the brain activity for the pre- and post-Drop events. In the first level analysis, multiple regression was used with subjective rating as a covariate. In the second level analysis, paired-sample t-tests were used to compare the modulated activity for pre- and post-Drop events.

## 3 Results

Three participants had to be excluded as their eye-blink activity was not within the first two components of spatial confounds. Therefore, data for 33 participants are reported here. Analysis of the 3D source reconstructed data showed significant increase in inferior and middle frontal gyri (IFG and MFG) activity, and significant decrease in pre and postcentral gyri (PreCG and PostCG) and precuneus (PCUN) activity in post-Drop, respectively. For a table summarising the analysis see *Table **1*** and also Figures 1-2.

**Figure 1.**
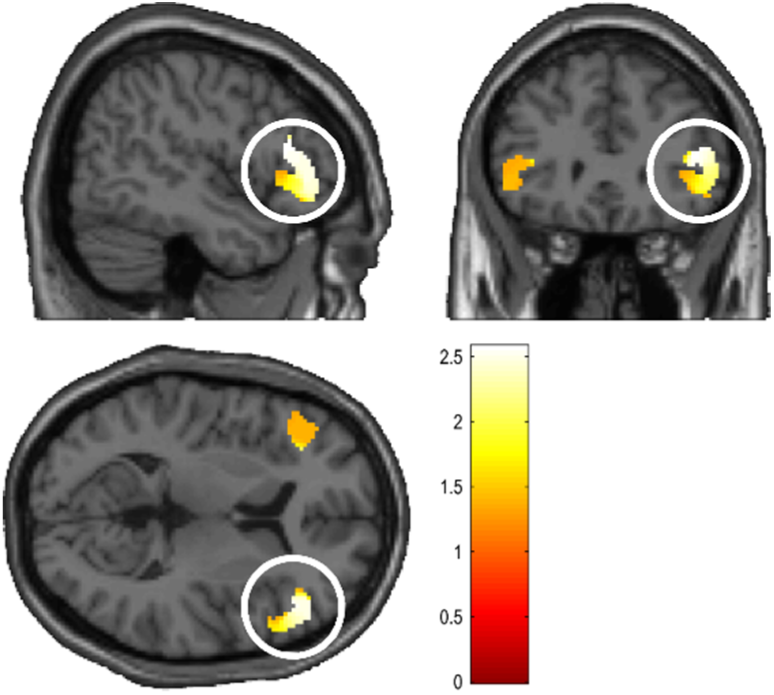
Main effect of Post > Pre showing the right inferior and middle frontal cortices (y = 30, z = 8) (*k* > 5 voxels, *P_FWE_* < 0.05).

**Figure 2.**
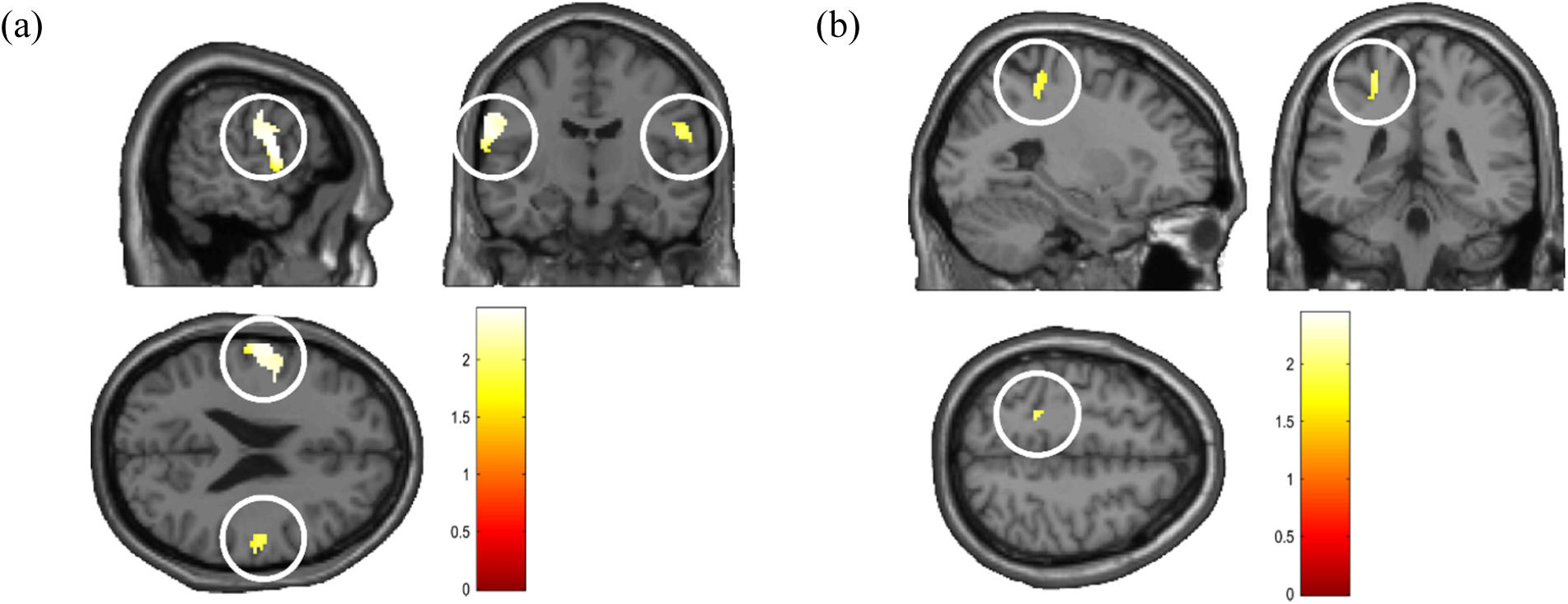
Main effect of Pre > Post (*k* > 5 voxels, *P_FWE_* < 0.05) (a) bilateral Postcentral Gyrus and the left Precentral Gyrus (y =11, z = 23), (b) left Precuneus (y = −40, z = 52).

**Table 1.** Summary of the analysis of the 3D source reconstruction of the EEG data.

|  |  |  |  |  |  |  |  | MNI |  |  |
| --- | --- | --- | --- | --- | --- | --- | --- | --- | --- | --- |
| | Brain Area | BA | $K_E$ | $P_{FWE}$ | $T$ | $Z$ | $P_{uncorr}$ | x | y | z |
| **Main Effect** |  |  |  |  |  |  |  |  |  |  |
| Post > Pre | r-Inf Frontal Gyrus | 9 | 1021 | 0.012 | 2.57 | 2.43 | 0.007 | 56 | 22 | 16 |
|  | r-Mid Frontal Gyrus | 46 |  | 0.015 | 2.47 | 2.35 | 0.009 | 46 | 38 | 10 |
| Pre > Post | l-Precentral Gyrus | 4 | 876 | 0.016 | 2.44 | 2.32 | 0.010 | -58 | 0 | 14 |
|  | l-Precentral Gyrus | 4 |  | 0.017 | 2.41 | 2.29 | 0.011 | -58 | -10 | 36 |
|  | l-Postcentral Gyrus | 3 |  | 0.019 | 2.37 | 2.26 | 0.012 | -64 | -12 | 28 |
|  | r-Postcentral Gyrus | 3 | 233 | 0.031 | 2.11 | 2.03 | 0.021 | 60 | -14 | 24 |
|  | l-Postcentral Gyrus | 40 | 244 | 0.039 | 2.00 | 1.93 | 0.027 | -26 | -38 | 60 |
|  | l-Precuneus | 7 |  | 0.041 | 1.98 | 1.91 | 0.028 | -24 | -40 | 52 |
| **Parametric Modulation** |  |  |  |  |  |  |  |  |  |  |
| Post > Pre | r-Inf Frontal Gyrus | 47 | 457 | 0.013 | 2.46 | 2.34 | 0.010 | 46 | 32 | -14 |
|  | r-Mid Frontal Gyrus | 46 |  | 0.016 | 2.38 | 2.27 | 0.012 | 52 | 26 | 20 |
|  | l-Inf Frontal Gyrus | 47 | 922 | 0.014 | 2.44 | 2.32 | 0.010 | -40 | 34 | -16 |

*Note:* BA: Brodmann Area, Inf: Inferior, Mid: Middle, FWE is family-wise error correction for a 3mm sphere around the peak voxel.

Parametric modulation analysis demonstrated significantly greater activity in bilateral IFG and MFG areas during post-Drop moments, meaning these areas correlate positively with higher subjective ratings of excitement in post-Drops (see Table 1 and Figure 3).

**Figure 3.**
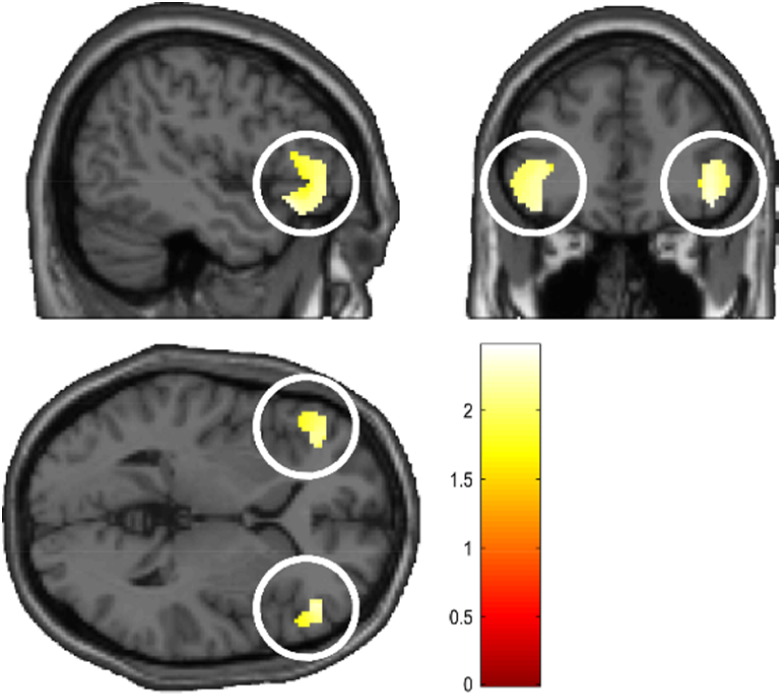
Parametric modulation of Post > Pre with subjective rating of excitement showing bilateral inferior and middle frontal cortices (y = 38, z = 2) (*k* > 5 voxels, *P_FWE_* < 0.05).

Further analysis showed a high correlation between the subjective strength of the Drop (as measured by pre-study questionnaire) and subjective rating of excitement during the study (r(88) = .60, p < 0.001).

## 4 Discussion

Here we offer the first exploration into the way music including a Drop relates to brain activity and high arousing positive emotions. We show that the structure of Drops in music strongly corresponds to significant brain activity across several regions, including; the bilateral inferior frontal gyrus (IFG; BA47), right middle frontal gyrus (MFG; BA46), bilateral pre- and post-central gyri (PreCG and PostCG; BA4 and 3), and precuneus (PCUN; BA7). Despite active areas not all being predicted, the current research suggests their association with music processing and emotions support our hypothesis. Also, IFG and MFG activity were moderated through ratings of excitement, implying these regions are associated with high arousal and positive emotions during Drop experiences. This suggests Drops can elicit strong and arousing emotions, alongside relevant neurological responses with positive valence.

Numerous brain regions, including all six here, have been associated with processing music and musical features. For instance, their activity have been related to passively, explicitly, and implicitly apprehending music over other auditory stimuli or silence (Altenmüller, Siggel, Mohammadi, Samii, & Münte, 2014; Bianco et al., 2016; Bogert et al., 2016; Plailly, Tillmann, & Royet, 2007; Proverbio & De Benedetto, 2018; Seger et al., 2013; Schön et al., 2010; Tabei, 2015). Also, these regions associate with perceiving rhythm, meter, melody, pitch, and tempo (Schön et al., 2010; Thaut et al., 2014). This suggests activity in these areas may relate to musical feature processing as they build and deviate over Drops duration.

### 4.1 Brain and Emotional Responses to Pre-Drop

Difference in brain activity between pre- and post-Drop suggests the regions’ functionality in music and emotional processing. For instance, the PreCG, PostCG, and PCUN were more active pre-Drop. Despite the PreCG and PostCG mostly being linked to motor functions, they have also been implicated in music processing and assimilating motor, somatosensory and auditory information (Potes, Gunduz, Brunner, & Schalk, 2012; Ramos-Murguialday & Birbaumer, 2015; Tanaka & Kirino, 2018). The PreCG, PostCG, and PCUN play a role in music attention, tempo alterations, absolute pitch, rhythm perception, and processing music intensity (Dohn et al., 2015; Ono, Nakamura, & Maess, 2015; Potes et al., 2012; Schön et al., 2010; Thaut et al., 2014). Thus, in support of our hypothesis, greater PreCG, PostCG and PCUN activity during pre-Drop listening could be associated with processing specific music features, including tempo, rhythm, and intensity as they build in aspects such as speed and volume to then stop or slow.

Also, these brain regions during pre-Drop moments are implicated in the processing of music-induced emotions. For example, PreCG and PostCG are linked to experiencing tension, expectancy, and anticipation (Bjork & Hommer, 2007; James, Michel, Britz, Vuilleumier, & Hauert, 2012; Lehne et al., 2014; Thaut et al., 2014; Vuilleumier & Trost, 2015). Also, the PCUN is used in evaluating music’s continuous structure, making perception-based predictions, and entrainment (Alluri et al., 2017; Trost et al., 2014). Thus, activity in the PreCG, PostCG and PCUN could be involved in processing musical structure and predicting future music events, alongside emotions such as tension, anticipation and expectancy. Greater activity pre-Drop may suggest listeners experience tension and anticipation as musical features build ahead of the predicted future deviation at post-Drop.

### 4.2 Brain and Emotional Responses to Post-Drop

IFG and MFG had greater activity post-Drops. It has been shown that the IFG contributes to perceiving music features (such as rhythm) and semantics (Lappe et al., 2013; Schön et al., 2010; Thaut et al., 2014). For example, greater IFG cortical thickness relates to possession of absolute pitch, and complex music, rhythm, and harmony apprehension (Dohn et al., 2015). This implies greater IFG activity post-Drop may index the processing of Drop features, as well as assessing familiarity of the Drops’ general structure as the music builds to then change.

Similar to the IFG, MFG activity also increased post-Drop and may relate to processing Drop features. For example, the MFG is particularly active within general and implicit music processing, working memory (WM; alongside the IFG), and music rule perception (Bogert et al., 2016). This implies the MFG’s importance in assessing Drops’ continuous musical structure through WM and structural processing, enabling predictions of post-Drop deviations and subsequent pre-Drop tension as listeners wait for the music changes to occur. Therefore, MFG activity (although not predicted) also supports the hypothesis that brain activity would increase due to the Drops’ specific structure of tension and deviation.

### 4.3 Correlational Brain Activity with Excitement

Importantly, only IFG and MFG activity were moderated by Drop excitement ratings, suggesting elevated activity post-Drop correlated with greater arousal and positive emotions as music deviates. Specifically, increased bilateral IFG and right MFG activity occurred post-Drop alongside greater excitement ratings, implying heightened activity related to experiencing high arousal during music deviation (Chapin, Jantzen, Kelso, Steinberg, & Large, 2010; Flores-Gutiérrez et al., 2007). For example, Flores-Gutiérrez et al. (2007) compared two classical works, one by Mahler (high arousal) and one by Bach (low arousal). They demonstrated that Mahler’s work was associated more with IFG and MFG activity. In support of our hypothesis, this implies the IFG and MFG are important to experiencing highly arousing emotions and may explain increased activity alongside greater excitement ratings. However, the IFG and MFG were also connected to highly arousing negative emotions, raising ambiguity as to whether they relate to highly arousing and positive emotions, such as excitement.

Nevertheless, the MFG and IFG are also connected with processing positive music-evoked emotions (Barrett & Janata, 2016; Kim et al., 2017; Wallmark et al., 2018). Greater MFG and IFG activity contributes to perceived, felt and pleasant music emotions; including nostalgia, happiness, liking, and empathy (Barrett & Janata, 2016; Brattico et al., 2011; Joucla et al., 2018; Kim et al., 2017; Koelsch et al., 2006; Tabei, 2015; Wallmark et al., 2018). Thus, greater MFG and IFG activity post-Drop and their correlation with excitement could suggest more positive emotive responses in listeners as music deviates (Bogert et al., 2016; Kohn et al., 2014; Wallmark et al., 2018). However, some pleasant emotions linked to IFG and MFG activity are lower in arousal, making it less clear whether IFG and MFG activity increases alongside greater positive emotions or arousal.

Currently, little research has linked the IFG and MFG to arousal in music, yet much research has associated these regions with positive emotions, including pleasantness (Altenmüller et al., 2014; Joucla et al., 2018). This suggests these regions may be more involved in processing positive emotions than high arousal and could be linked to excitement due to its positive valence, although more research is needed.

### 4.4 Linking Drop Structure, Brain Activity, and Emotions

One way to explain how Drops structure may contribute to brain activity, high arousal, and positive music emotions, is via Huron’s (2006) ITPRA theory. Specifically, musical tension and prediction are important to processing and experiencing music emotions (Huron, 2006; Koelsch, 2014). Music follows certain structures or rules within time, musical events, intensity and space, enabling musical syntax which is associated with IFG activity (Asano & Boeckx, 2015; Lehne et al., 2014; Koelsch, 2014; Kunert et al., 2015). Musical syntax facilitates predictability and listeners’ anticipation of future music moments, which relates to musical tension, positive emotions when predictions are correct, and co-occurring IFG activity (Bianco et al., 2016; Huron, 2006; Koelsch, 2014; Lehne et al., 2014; Schön et al., 2010; Zatorre, Chen, & Penhune, 2007). Thus, the ITPRA theory suggests music evokes emotions via tension and anticipation of predicted moments, and tension has been shown to relate to IFG activity.

Specifically, Drops possess a unique syntax and repeated listeners of dance music may learn to predict post-Drop deviations. However, there is uncertainty of when deviations will occur due to delays from pre-Drop moments. This pre-Drop delay, including rising musical features, may cause greater tension associating with enhanced PCUN activity (Vuilleumier & Trost, 2015). As musical features deviate post-Drop, this heightened tension may amplify positive emotions due to contrastive valence, as well as mediating IFG and MFG activity (Bianco et al., 2016; Lappe et al., 2013; Seger et al., 2013). Therefore, Drops’ tension-inducing and deviating structure may link to ITPRA via tension and prediction, which may also mediate activity in the PCUN pre-Drop and IFG and MFG post-Drop, as well as being associated with greater arousing and positive emotions, including excitement.

Also according to ITPRA, contrastive valence suggests Drops with longer and more tension inducing pre-Drops preceding predicted large deviations post-Drop, would evoke greater positive emotions, including excitement (Huron, 2006). Parametric modulation results showing greater IFG and MFG activity alongside increased excitement ratings, could suggest their importance in processing music predictions and tension, and to experiencing subsequent positive emotions (Lehne et al., 2014; Pecenka, Engel, & Keller, 2013). Thus, larger IFG and MFG activity post-Drop could relate to the processing of tension, music prediction and resulting positive emotion as expected post-Drop deviations occur.

However, our finding of no significant alterations in DLPFC activity across Drop apprehension contradicts the hypothesis that DLPFC activity will increase in relation to Drop processing. In addition to many other functions, the DLPFC is involved in music processing, specifically it is more active in processing familiarity, rhythm and sequenced sounds, as well as detecting pitch alterations or deviances (Altenmüller et al., 2014; Plailly et al., 2007; Flores-Gutiérrez et al., 2007; Seger et al., 2013; Platel et al., 1997; Koelsch & Siebel, 2005, Doeller et al., 2003; Dohn et al., 2015, respectively). Also, greater DLPFC activity is associated with likable music, increased physiological arousal, and pleasant emotions, including complex positive feelings such as empathy and nostalgia (Barrett & Janata, 2016; Bigliassi, León-Domínguez, & Altimari, 2015; Brattico et al., 2011; Joucla et al., 2018; Koelsch et al., 2006; Wallmark et al., 2018). Although DLPFC activity may be genre specific and is not always active during music apprehension (Bigliassi et al., 2015). For example, increased DLPFC activity is associated with classical music listening but not techno music (Bigliassi et al., 2015). Therefore, less DLPFC activity raises ambiguity as to its importance in processing modern electronic music and musical deviations in Drops, which may explain lack of DLPFC activity changes here. Future research should evaluate further the limited processing ability of the DLPFC in alternative music genres.

### 4.5 Conclusions

This paper offers a first insight into the neurological responses to music that contains a Drop. Activity in several brain areas including; the IFG, MFG, PCUN, and PreCG and PostCG during either pre- or post-Drop may relate to the processing of specific music features, tension, prediction, and emotions. These include pitch, rhythm, and deviation, as well as the elicitation of high arousal and powerful, positive emotions, such as excitement. Future research should expand on the relationship between Drops, brain activity and different emotional responses. It should attempt to establish a causal relation between the aforementioned brain regions, and emotional and arousing responses using stimulation and other imaging techniques. For instance, future research may compare neurological responses to Drops with other music genres, such as classical. Also, future research could use cathodal transcranial direct current stimulation (tDCS) within the IFG and MFG to assess music processing and emotional differences.

